# Single-cell analysis of developing B cells reveals dynamic gene expression networks that govern B cell development and transformation

**DOI:** 10.1101/2020.06.30.178301

**Authors:** Robin D. Lee, Sarah A. Munro, Todd P. Knutson, Rebecca S. LaRue, Lynn M. Heltemes-Harris, Michael A. Farrar

**Affiliations:** Department of Laboratory Medicine and Pathology, Center for Immunology, Masonic Cancer Center, University of Minnesota; Minnesota Supercomputing Institute, University of Minnesota

**Keywords:** B cell development, Single-cell RNA-sequencing, CITE-Seq, Hashtagging, pre-B cell expansion, EBF1, YBX3, pseudotime, B cell acute lymphoblastic leukemia (B-ALL)

## Abstract

Integration of external signals and B-lymphoid transcription factor activities orchestrate B cell lineage commitment through alternating cycles of proliferation and differentiation, producing a diverse repertoire of mature B cells. We used single-cell transcriptomics and proteomics to characterize B cell development. Our analysis revealed unique transcriptional signatures that refine the pre-B cell expansion stages into novel pre-BCR-dependent and pre-BCR-independent proliferative phases. These changes correlate with unexpected dynamic and reciprocal changes in expression of the transcription factor EBF1 and the RNA binding protein YBX3, that are defining features of the pre-BCR-dependent stage. Using pseudotime analysis, we further characterize the expression kinetics of different biological modalities across B cell development, including transcription factors, cytokines, chemokines, and their associated receptors. Our findings reveal the underlying heterogeneity of developing B cells and point to key developmental nodes linked to B cell transformation.

## Introduction

Distinct stages of B cell development have been delineated using flow cytometry and a variety of surface (Hardy et al., 1991; Rolink et al., 1994) and intracellular markers (Billips et al., 1995; Ghia et al., 1998). The use of such markers in combination with distinct gene knock-out mice has greatly expanded our understanding of the role that specific B-lymphoid transcription factors (Nutt et al., 1999; Schwickert et al., 2014; Sö ren Boller et al., 2016), cytokines (Corcoran et al., 1998; Goetz et al., 2004; Mandal et al., 2019), and signaling pathways play in entraining B cell development. However, these markers are insufficient to fully demarcate distinct subsets (Hardy et al., 1991), resulting in the analysis of mixed populations. These limitations have led to an incomplete understanding of B-lymphoid transcription factor expression kinetics across the B cell developmental trajectory, and the orchestration of transcriptional programs underlying the alternating cycles of proliferation and differentiation. Capturing transition states as B cells differentiate from one stage to the next is particularly difficult. Furthermore, perturbations during normal B cell differentiation can lead to development of B-cell acute lymphoblastic leukemia (B-ALL) (Gu et al., 2019; Heltemes-Harris et al., 2011; Mullighan et al., 2007). However, exactly what stages are most permissive for transformation remains imprecisely defined. Recent characterization of B-ALL subtypes showed diverse transcriptional signatures, suggesting multiple points of origin, or use of different signaling pathways to transform (Gu et al., 2019; J. F. Li et al., 2018). Therefore, understanding normal B cell transcriptional programs can determine where transformation occurs and how B-ALLs exploit B cell developmental pathways.

To address the above questions, we used single-cell transcriptomics (scRNA-Seq) and proteomics (CITE-Seq; <u>C</u>ellular <u>I</u>ndexing of <u>T</u>ranscriptomes and <u>E</u>pitopes by <u>Seq</u>uencing) (Stoeckius et al., 2017) to precisely characterize different subsets of B cell development. Our analysis discovered several novel stages of pre-B cell differentiation – including a pre-BCR-dependent and two pre-BCR-independent stages that exhibited distinct modalities of proliferation. This process of pre-B cell differentiation was reflected by unexpected dynamic regulation of the transcription factor EBF1, downstream of the pre-BCR, and the RNA binding protein YBX3. In contrast, the pre-BCR-independent stages correlated with changes in chemokine and cytokine receptors and suggest that these stages may involve differential localization of pre-BCR-independent stage subsets within the bone marrow. Finally, comparisons of various human B-ALL transcriptomes to those of different stages of B cell development highlight the pre-BCR-dependent proliferation stage as a critical node that B-ALL subtypes hijack during transformation.

## Results

### Identifying B cell development stages using scRNAseq and CITE-Seq

To couple transcriptional information with B cell stage-defining surface marker expressions, we used combined single-cell RNA sequencing and CITE-Seq (also referred as <u>A</u>ntibody-<u>D</u>erived-<u>T</u>ags [ADT] hereafter) proteomics. Bone marrow from two wildtype C57BL/6 mice were harvested and stained with two distinct oligo-labeled antibodies that recognize CD45 and MHC class I, which allow identification of cells derived from each individual mouse (referred to as hashtag antibodies). We further stained cells with a panel of CITE-Seq antibodies (B220, CD19, CD93, CD25, IgM, and CD43) as well as fluorescently labeled B220 and CD43. Cells were sorted at a 1:1 ratio of B220^+^CD43^+^ and B220^+^CD43^−^ cells to enrich for early progenitor B cell subsets (Figure 1A). After processing samples, the data set contained 7454 single cells contributing to 14 transcriptionally unique clusters (Figure 1B). Using a list of previously described cell cycle genes (Nestorowa et al., 2016), we labeled cells for their cell cycle status and found that the proliferating cells (S and G2/M phase) were clustered together away from quiescent cells (G1), suggesting that cycling status was a major source of variance (Figure 1C). Consistent with this observation, *Mki67* gene expression levels were high in cells classified as S or G2/M phase (Figure 1C). Further, we found that CITE-seq provided superior sensitivity compared to the corresponding transcript expression (Figure 1D). Cells marked CD43^+^ by CITE-Seq comprised the majority of cycling cells, supporting previous characterizations of cycling progenitor B cells (Hardy et al., 1991). Both wildtype mice were equally represented in all cell clusters (Supplementary Figure 1A) and had equal detection of all CITE-seq antibodies (Supplementary Figure 1B and 1C). Overall, CITE-seq antibody expression recapitulated flow cytometry-based staging of B cell development and when coupled with transcriptomic signatures provided the basis for demarcating different stages and transitions during B cell development.

**Figure 1.**
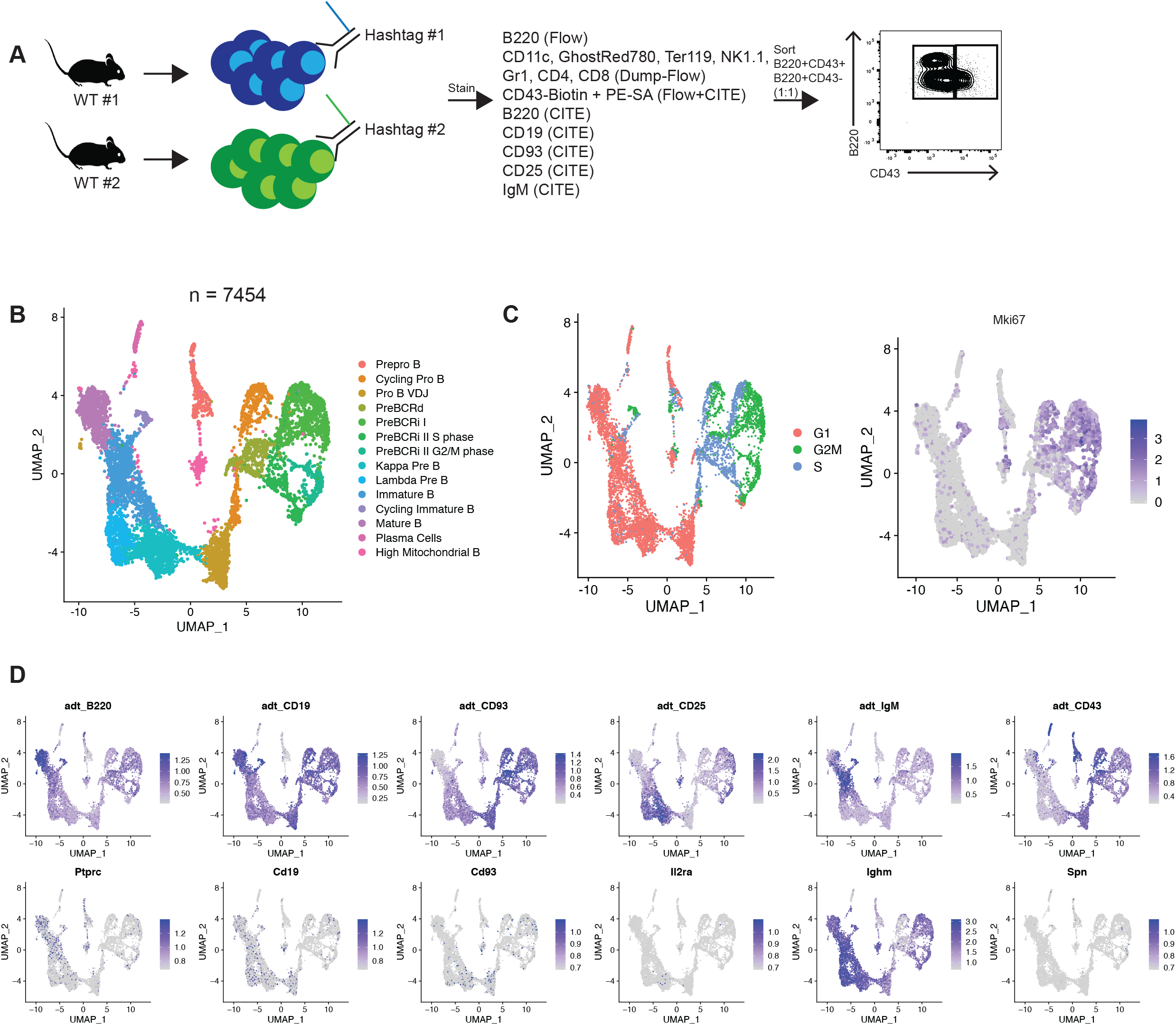
Identifying B cell development stages using scRNAseq and CITE-Seq. **A.** Schematic of experimental setup. **B.** UMAP dimension reduction projection of all cells (n = 7454) from two wildtype C57BL6 mice. Fourteen clusters were identified and corresponding population names for each cluster are listed (Pre-BCRd = Pre-BCR-dependent, Pre-BCRi = Pre-BCR-independent) **C.** Feature plot of cells that are labeled according to their cell cycle status based on gene expression (left) and *Mki67* transcription expression (right). **D.** Feature plot of cells for their CITE-Seq/ADT antibody expression (top row) and corresponding gene transcript expression (bottom row).

### Transcriptional signatures of pre-pro B cells and committed pro B cells

We first used antibody derived CITE-Seq tags to define B220^+^CD43^+^CD19^−^ pre-pro B cells. These cells expressed early B-lineage associated genes such as *Flt3*, *Il7r* and *Cd79a* (Figure 2A). Pre-pro B cells have also been shown to express genes associated with myeloid lineages, consistent with the observation that they can give rise to myeloid cells as well (Hystad et al., 2007; Rumfelt et al., 2006). Indeed, we found that the pre-pro B cells have high expression of myeloid lineage-associated transcription factors such as *Runx2*, *Irf8*, and *Tcf4,* and plasmacytoid dendritic cells markers, such as *Bst2;* these genes were silenced upon commitment to the B cell lineage at the pro-B cell stage (Figure 2B). We also assessed the expression of previously described EBF1-repressed target genes, including *Tyrobp*, C*lec12a*, *Cd300a*, *Cd7*, *Chdh* and *Mycl* (R. Li et al., 2018). (Figure 2C). These target genes were highly expressed in the pre-pro B cell cluster, while *Ebf1*-expressing pro-B cells had low or undetectable expression of these genes, further distinguishing the pre-pro B cells from pro-B cells (Figure 2B). Pro-B cells, traditionally delineated as c-KIT^+^ cells (Ogawa et al., 2000) had *Kit* gene expression (Figure 2D). In addition, pro-B cells expressed the *Bcl-2* family gene, *Bok,* and interferon-stimulated genes, such as *Ifitm2* and *Ifitm3* (Figure 2D, Supplementary Table 1). Finally, EBF1 positively regulates the expression of pre-BCR surrogate light chain genes, *Vpreb1* and *Igll1* (Sigvardsson et al., 1997). We found that cycling pro-B cells had high expression of both *Vpreb1* and *Igll1*, compared to the pre-pro B cells that do not express *Ebf1* (Figure 2E). Thus, our analysis confirmed several genes expression patterns associated with the transition from pre-pro-B to pro-B cells, as well as identified novel highly specific markers of pro-B cells such as *Bok*.

**Figure 2.**
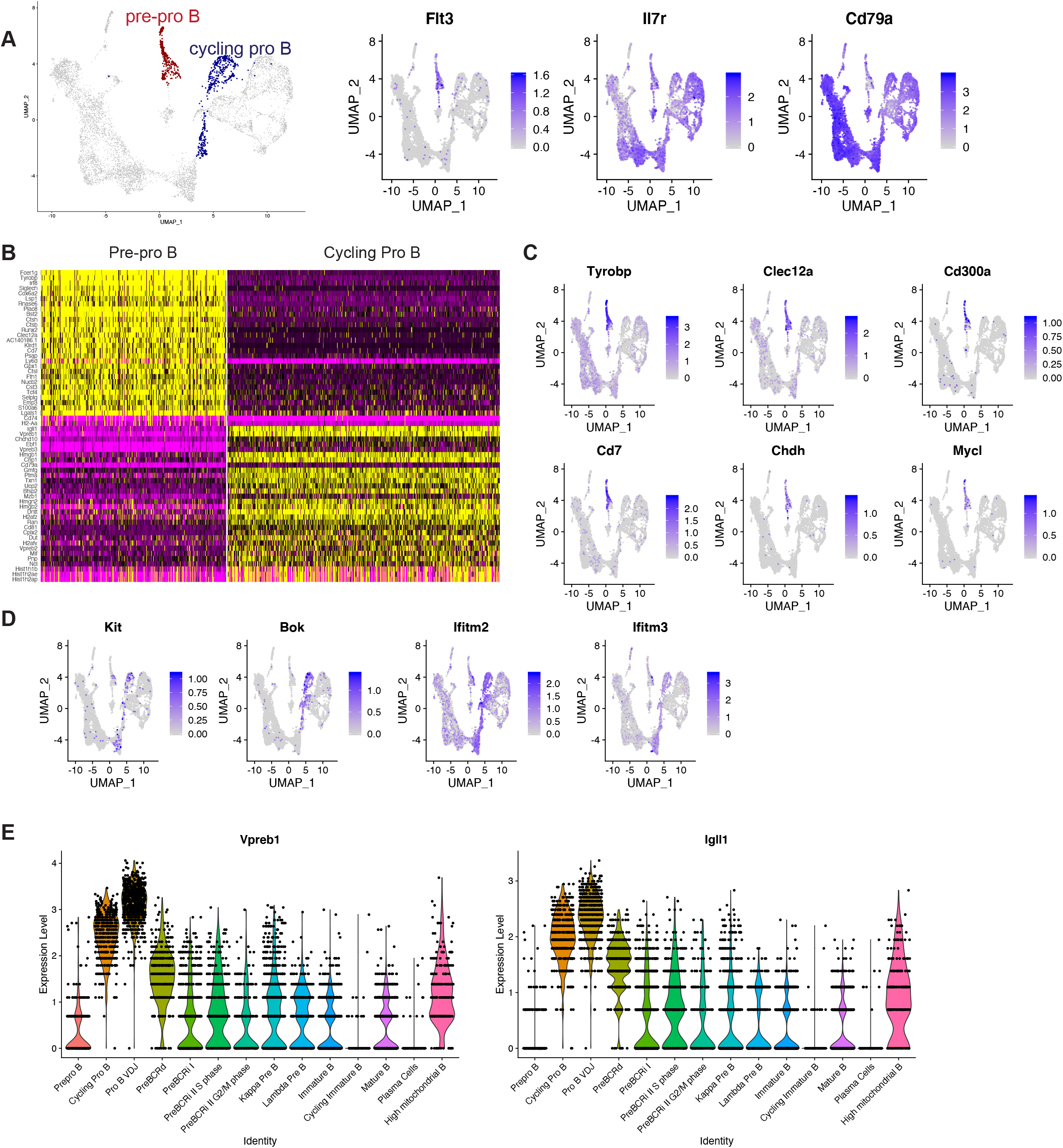
Transcriptional signatures of pre-pro B cells and committed pro B cells. **A.** Highlighted populations for pre-pro B and cycling pro B cells (left). Feature plots for early B cell markers (right). **B.** Heatmap of genes that are differentially expressed between the pre-pro B cells and the cycling pro B cells. **C.** Feature plot of EBF1-target genes **D.** Feature plot of genes denoting the pro B cell stage (*Kit*), or genes uniquely expressed during pro B cells. **E.** Violin plot of the expression of surrogate light chains, *Vpreb1* and *Igll1* across the B cell development stages.

### Pre-B cell expansion is comprised of distinct Pre-BCR-dependent and Pre-BCR-independent proliferation stages

Next, we assessed the transcriptional signature of the ADT-B220^+^CD19^+^CD43^+^CD25^−^ cells. We refer to these cells as the Pre-BCR-dependent proliferation cluster (also referred to pre-BCRd hereafter) (Figure 3A). Pre-BCR signaling initiates silencing of the surrogate light chain locus(Parker et al., 2005). In accordance with this, the Pre-BCR-dependent cells have intermediate expression of both *Vpreb1* and *Igll1* (Figure 2E). Genes that were uniquely upregulated in this cluster included *Nrgn* and *Ybx3* (Figure 3A, Supplementary Figure 2A and Supplementary Table 2). NRGN (Neurogranin) is a calmodulin-binding protein that regulates the dynamics of calcium binding to calmodulin (Hoffman et al., 2014). Neurogranin is upregulated in activated (Glynne et al., 2000) and anergic B cells (Merrell et al., 2006), which suggests that neurogranin expression is controlled by B cell receptor signaling. YBX3 is a DNA/RNA binding protein that was recently shown to stabilize the amino acid transporter transcripts *Slc7a5* and *Slc3a2* in HELA cells, allowing for their robust translation (Cooke et al., 2019). SLC7A5 and SLC3A2 heterodimerize to form CD98, a large neutral amino acid transporter. Expression of CD98 in CD8^+^ T cells has been shown to be tightly controlled by antigen receptor signaling and is critical for MYC expression (Sinclair et al., 2013). Our findings suggest that pre-BCR signaling may serve a similar function. Consistent with this idea, *Myc* was most highly expressed in the *Ybx3*, *Slc7a5*, *Slc3a2* expressing pre-BCRd cluster (Figure 3A). Surprisingly *Ebf1* expression is significantly reduced, while expression of the transcription factor *Pax5* was largely unchanged and *Ikzf1* was modestly induced (Figure 3B). Concordant with low *Ebf1* expression, EBF1-target genes, such as *Cd79a* (Hagman et al., 1991; Sigvardsson et al., 2002) and *Cd79b* (Åkerblad et al., 1999), are also significantly reduced in the pre-BCRd stage (Figure 3C). In addition, *Il7r* expression, a negative regulator of pre-BCR signaling components (Katerndahl et al., 2017), is also reduced (Figure 3C). To identify potential transcriptional regulators that govern this pre-BCR-dependent expansion stage, we performed a Landscape In Silico deletion Analysis (LISA) (Qin et al., 2020) using the top 100 differentially upregulated genes in the pre-BCRd cluster. After excluding factors not expressed in B cells (i.e. MYCN), we found MYC to be the top predicted regulator of this gene set in pre-BCRd cells. Consistent with its decreased expression, EBF1 was predicted to have minimal contributions in Pre-BCRd cells (Figure 3D), suggesting that MYC is the critical transcription factor that governs the transcriptional landscape during Pre-BCR-dependent expansion. To further understand the importance of *Ebf1* downregulation during pre-BCR-signaling, we examined differentially regulated genes that contained EBF1 binding sites. EBF1 has clear binding sites at the promoters for *Nrgn* and *Myc* (Figure 3E). Likewise, EBF1 binds to a known superenhancer linked to the *Myc* locus (Katerndahl et al., 2017). In addition, potential binding sites were observed within the promoters of the *Slc7a5* and *Slc3a2* genes (Figure 3E). To determine how EBF1 affects expression of these genes, we performed RNA-seq in wildtype and *Pax5*^*+/−*^ x *Ebf1*^*+/−*^ leukemic progenitor B cells. We found that *Slc7a5* and *Slc3a2*, along with other pre-BCR-dependent stage module genes (*Ybx3*), were upregulated with decreased *Ebf1* gene dosage (Figure 3F). This is consistent with our recent observation that *Myc* expression is increased in *Pax5^+/−^ x Ebf1^+/−^* pre-leukemic and *Pax5^+/−^ x Ebf1^+/−^* leukemic progenitor B cells (LMHH and MAF, submitted). These findings suggest that EBF1 mediates repression of the pre-BCR gene expression module.

**Figure 3.**
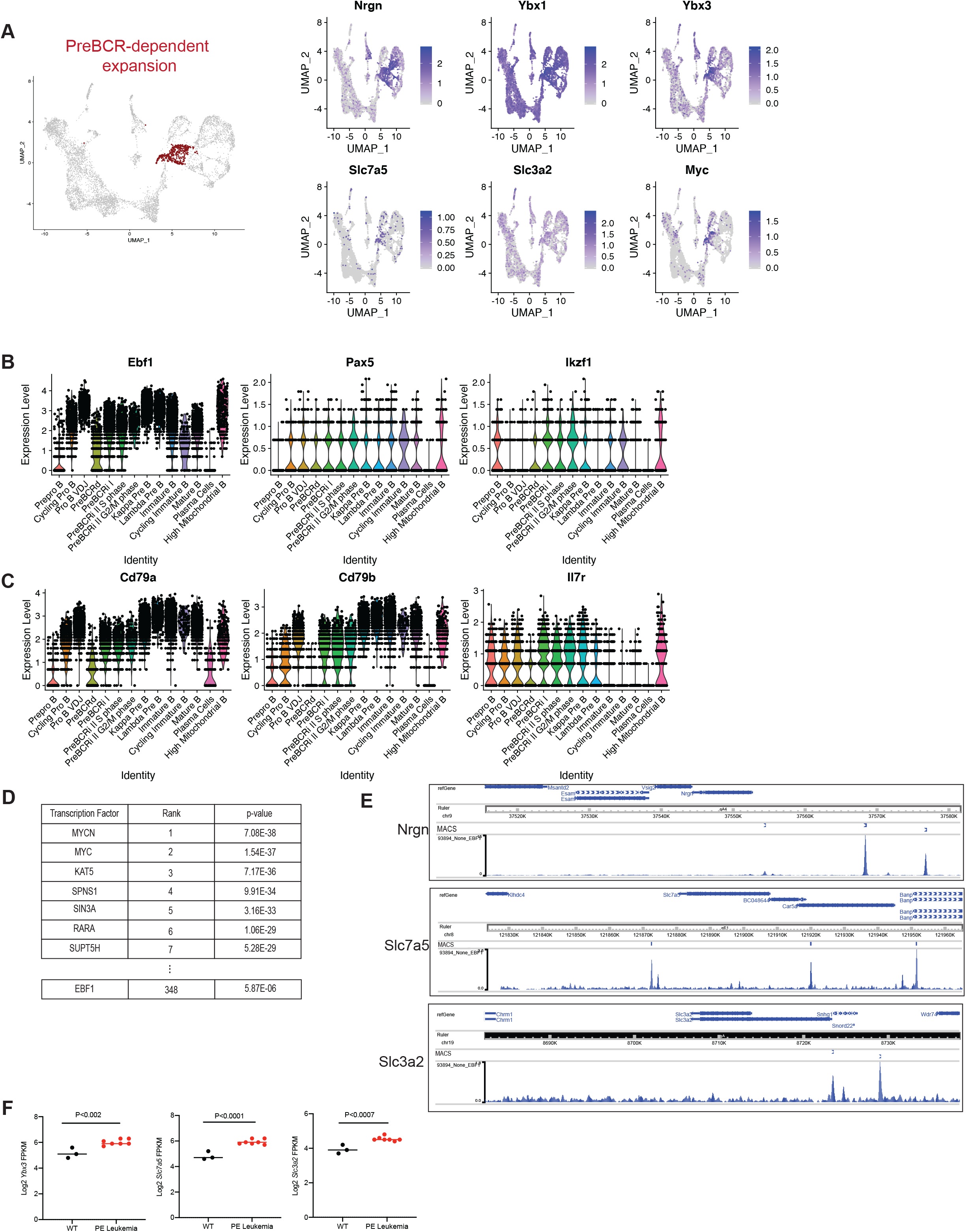
The transcriptome of the Pre-BCR-dependent expansion stage is governed by MYC-associated gene expression networks and requires repression of *Ebf1* expression. **A.** Highlighted population of the pre-BCR-dependent cluster (left) and feature plots for the pre-BCR-dependent markers highly expressed during this stage (right). **B.** Violin plot of B-lymphoid transcription factor expression, including *Ebf1*, *Pax5* and *Ikzf1* across B cell development. **C.** Violin plot of additional genes that are downregulated during pre-BCR-dependent proliferation, which includes *Il7r*, and EBF1-target genes such as *Cd79a*, *Cd79b*. **D.** Landscape *In silico* Deletion Analysis (LISA) to predict the transcriptional regulator of the top 100 differentially upregulated genes during pre-BCR-dependent proliferation. **E.** EBF1 binding sites at the promoters of *Nrgn*, *Slc7a5* and *Slc3a2* with indicated MACS peak calls(R. Li et al., 2018). **F.** Log2 transformed FPKM expression values obtained from RNA-seq of wildtype and *Pax5^+/−^ x Ebf1^+/−^* leukemic progenitor B cells. Progenitor B cells were obtained via negative selection of CD11c, TER119, GR1, Ig Kappa, Ig Lambda and positive selection of CD19. Statistical significance was determined using an unpaired student t-test.

YBX3 binds to *Jak1* transcripts and inhibits translation of *Jak1* in HELA cells (Cooke et al., 2019). To determine if YBX3 levels correlate with reduced JAK1 in pre-B cells, and to validate EBF1 downregulation during pre-BCR signaling, we used flow cytometry to characterize B220^+^CD19^+^CD43^+^CD98^high^ expressing cells. Ki67^+^JAK1^low^ cells have lower EBF1 expression (which correlates with high *Ybx3*) compared to the Ki67^+^JAK1^high^ cells (Supplementary Figure 3A). When evaluating the expression of Y-box family genes, *Ybx1* was ubiquitously expressed through all developmental stages, with peak expression during the pre-BCRd stage, whereas *Ybx3* expression was selectively induced during the pre-BCRd stage (Supplementary Figure 2A). We assessed the functional role of YBX3 during B cell development in *Ybx3*^*−/−*^ mice using flow cytometry. *Ybx3*^*−/−*^ B lineage cells had no significant phenotypic defects during the early proliferative phases (Hardy Fractions A-C; Supplementary Figure 3B, 3D) and late stage differentiation of B cell development (Hardy Fractions D-F; Supplementary Figure 3C, 3D). This is consistent with previous findings that YBX1 and YBX3 have redundancies in both function (Lu et al., 2006) and target mRNA binding (Lyabin et al., 2020). Collectively, our data highlights the reciprocal regulation of *Myc* and *Ebf1* during B cell development, where MYC is critical for governing differentially expressed genes in the pre-BCRd cluster. Furthermore, pre-BCR-signaling limits the IL7R-signaling axis by downregulating *Il7r* expression and JAK1 protein translation.

We next examined the identity and signature of the remaining cycling cells (Figure 4A). These cells have minimal surrogate light chain expression (Figure 2E), indicating further silencing of the locus. Notably, these cells have intermediate expression of the ADT-CD43 (Figure 4A), which suggests that they are transitioning towards the quiescent CD43^lo/−^ small pre-B cell stage. Some cells within these clusters also express CD25 protein (Figure 4A), a marker for pre-B cells (Rolink et al., 1994). A subset of these cells still expressed high levels of *Nrgn* (Figure 3A), which may suggest continued pre-BCR signaling. However, unlike the pre-BCR-dependent cluster, these cells have high expression of *Bach2* (Figure 4E), a transcription factor that restrains antigen receptor signaling (Roychoudhuri et al., 2016; Sidwell et al., 2020), and low expression of *Ybx3* and *Myc* (Figure 3A and 4B). This suggests that these pre-B cells are proliferating independently of pre-BCR signaling. When comparing the differentially expressed genes between the pre-BCR-dependent cells and the pre-BCR-independent cells from cluster I (Pre-BCRi I; Figure 4A), we found that pre-BCR-independent cells re-express high levels of *Il7r* and *Ebf1* (Figure 3B, 3C, and 4C), in accordance with restrained pre-BCR signaling. When comparing the two different pre-BCR-independent clusters (pre-BCRi I and preBCRi II), pre-BCRi I had heightened expression of histone genes (*Hist1h2ae*, *Hist1h2ap*, *Hist1h1b*, *Hist1h1e*, and *Hist1h4d*) (Figure 4D), but no difference in *Mki67* expression (Supplementary Figure 2B).

**Figure 4.**
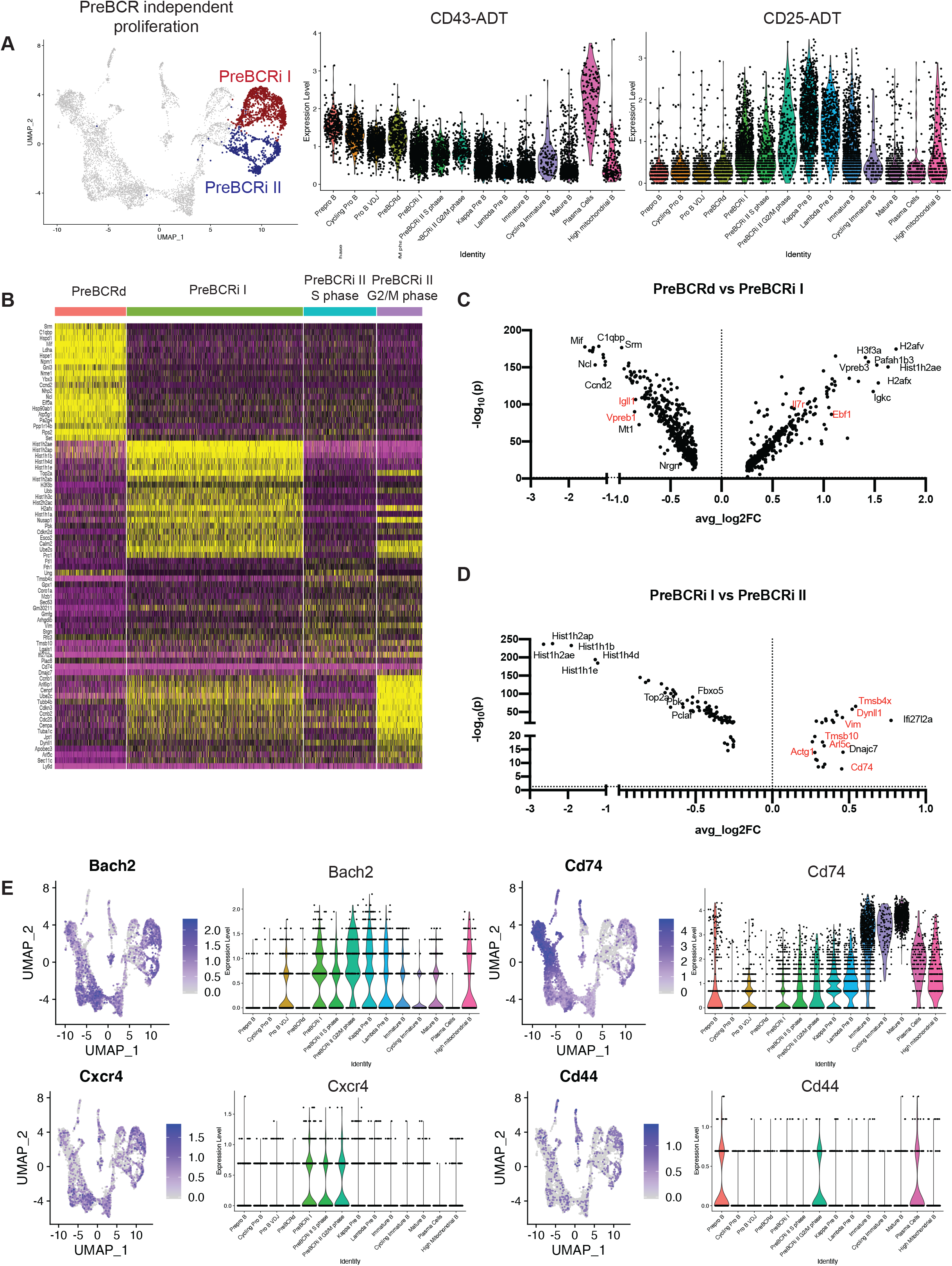
Pre-BCR-independent proliferation is distinct from pre-BCR-dependent proliferation. **A.** Highlighted Pre-BCRi I and Pre-BCRi II populations (left). Violin Plots of expression of ADT-CD43 and ADT-CD25 across the B cell development stages (right). **B.** Heatmap of differentially expressed genes between the Pre-BCR-dependent (Pre-BCRd), Pre-BCR-independent I (Pre-BCRi I) and Pre-BCR-independent II S, or G2/M phase. **C.** Volcano plot showing differentially regulated genes between the pre-BCRd cluster and the pre-BCRi I cluster. **D.** Differentially regulated genes between the pre-BCRi I cluster and the pre-BCRi II cluster. **E.** Single-cell RNA seq expression for *Mif* among stromal cells in the bone marrow microenvironment taken from data by Ref.(Tikhonova et al., 2019). Fbn1^high^Igf1^high^ osteogenic cell (O2) have the highest expression of *Mif*. **F.** Feature plot and violin plot for *Cd74*, *Cd44* and *Bach2*.

On the contrary, the Pre-BCRi II cluster is enriched for cells expressing cell motility-associated actin (*Actg1*), dynein (*Dynll1*) and thymosin (*Tmsb4x* and *Tmsb10*) genes (Figure 4D). Interestingly, expression of *Cxcr4* is further induced as pre-B cells reach the pre-BCRi stages (Figure 4E). Expression of CXCR4 would promote targeting of pre-BCRi cells to CXCL12-expressing cells, which also often express high levels of IL7 in the bone marrow (Cordeiro Gomes et al., 2016; Tikhonova et al., 2019). Since *Il7r* also increases in pre-BCRi cells these changes in gene expression suggest that pre-BCRd cells rely on pre-BCR signals for survival/ proliferation while pre-BCRi cells rely on IL7 signals. Although the IL7R and CXCR4 likely play the predominant role in promoting pre-BCRi cell survival, we did observe that pre-BCRi cells also started to express *Cd74* as well as *Cd44* (Figure 4E), which complex together to form a receptor for the chemokine ligand, Macrophage Inhibitory Factor (MIF). This could promote chemotaxis of pre-BCRi cells to *Mif* expressing Fbn1^high^Igf1^high^ osteogenic cells (Tikhonova et al., 2019),(Klasen et al., 2014) that also express IGF1 (Tikhonova et al., 2019), a paracrine growth factor that has been previously shown to be important for the generation of small pre-B cells (Yu et al., 2016). Thus, pre-BCRi cells likely require CXCR4/IL7R signals, with a possible contribution from the MIF/CD74/CD44 and IGF1/IGF1R signaling axes, to efficiently transit from a pre-BCR-dependent to a pre-BCR independent state.

### B cell differentiation and maturation

Quiescent IgM^−^ cells have high expression of *Rag1* and *Rag2*, suggesting V(D)J recombination of pro-B and pre-B cells (Figure 5A). While *Igkc* expression is detectable in all subsets except for cycling pro B cells and pre-BCR-dependent B cells, *Iglc1*, *Iglc2*, and *Iglc3* are only expressed in a subset of cells (Figure 5A). Furthermore, different B cell stages exhibited differential expression of trace-element-associated genes such as selenoprotein genes. *Selenom* was expressed in a subset of pro B VDJ cells (Supplementary Figure 4A). *Selenop* was expressed in both pro B and pre B cells undergoing recombination, whereas *Selenoh* was highly expressed in cycling pro B and pre B cells (Supplementary Figure 4A). The significance of this differential gene expression program remains to be ascertained, although selenium has been associated with immune function and activation (Hoffmann and Berry, 2008). Finally, the IgM^+^ cells were broken down into three clusters corresponding to immature B cells, cycling immature B cells and mature B cells. The cycling immature B cells have high expression of surface IgM. (Supplementary Figure 4B). Immature B cells express *Ms4a1* (CD20), whereas the mature B cells express *Ms4a4c*, *H2-Aa*, *Sell* (L-selectin), and *Ltb* (Figure 5B). Notably, a subset of mature B cells expressed *Apoe* (Figure 5B) and showed overlapping detection of both IgM and CD43 (Figure 1D). This subset shares features with previously described B1 bone marrow B cells (Choi et al., 2012), although the role of *Apoe* in these cells remains unknown.

**Figure 5.**
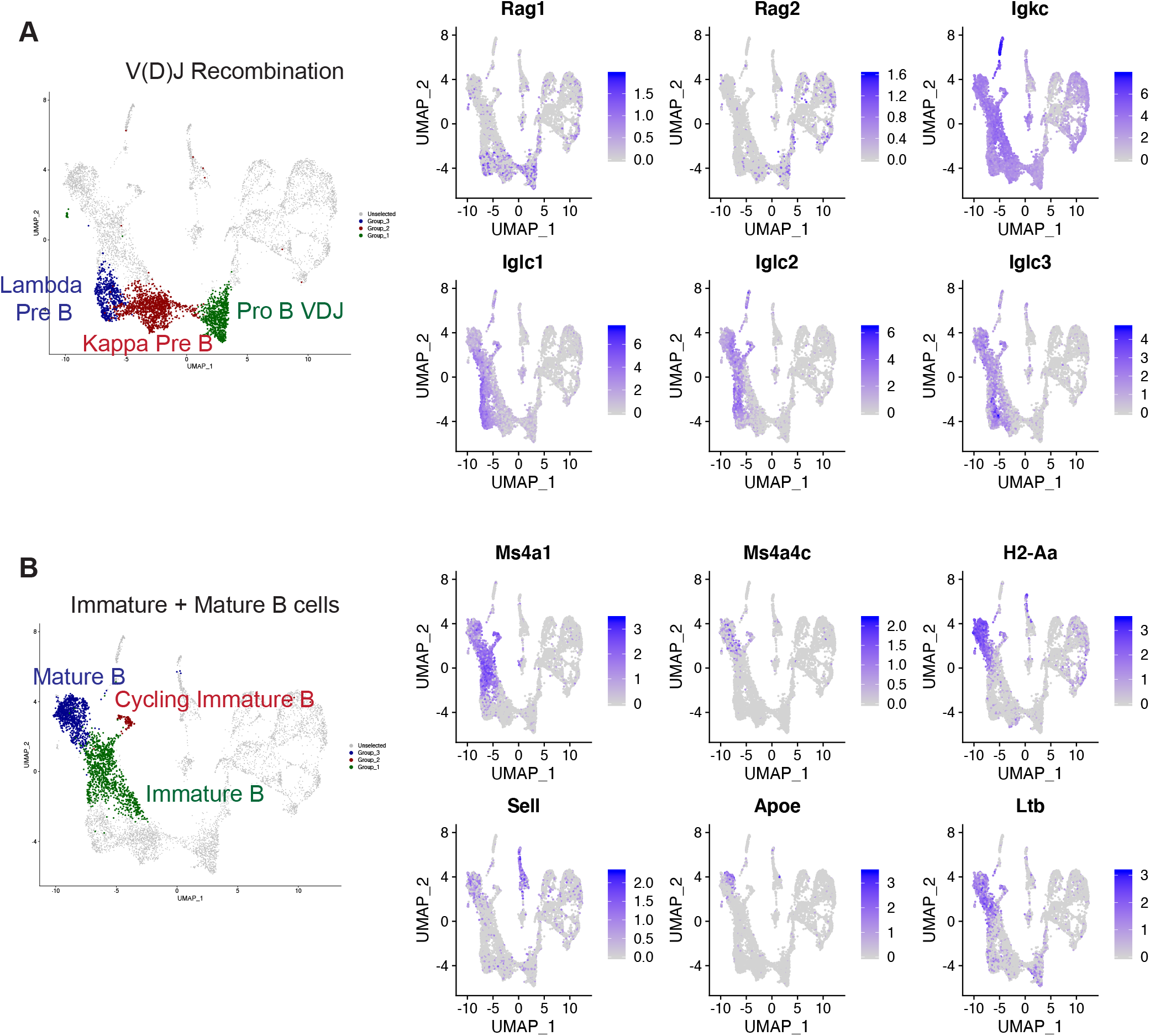
B cell differentiation and maturation. **A.** Highlighted B cell clusters undergoing V(D)J recombination (left). Feature plot of genes involved in V(D)J recombination or specific for pre B cell expression (right). **B.** Highlighted B cell clusters for late B cell maturation (left). Feature plot of genes highly expressed in immature or mature B cells (right)

### Pseudotime and module analysis of the B cell development trajectory

To understand the relationship between developmental stages and changes in gene expression over the B cell developmental trajectory, we performed pseudotime analysis using Monocle (Trapnell et al., 2014). To lessen the influence of cell cycle on UMAP positioning, we regressed out cell cycle genes within Monocle and performed UMAP dimension reduction. Using stage-defining markers, we identified 13 distinct B cell developmental stages (Figure 6A and Supplementary Figure 5) and calculated their respective pseudotime values to establish a developmental trajectory (Figure 6B). This analysis let us identify modules of genes that are changing across the developmental trajectory (Figure 6C). Gene ontology analysis of these gene modules revealed that the early B cell stages, including pre-pro B and pro-B cells, are significantly enriched for cell adhesion processes, whereas these signals are diminished in pre-B cells (Module 1 and Module 8; Figure 6C and 6D). This is in accordance with previous findings that pro B cells strongly promote adhesion to IL7-producing stromal cells (Fistonich et al., 2018), whereas pre-BCR-signaling and its downstream target IKZF1, are important for downregulating stromal adhesion components (Joshi et al., 2014; Schwickert et al., 2014).

**Figure 6.**
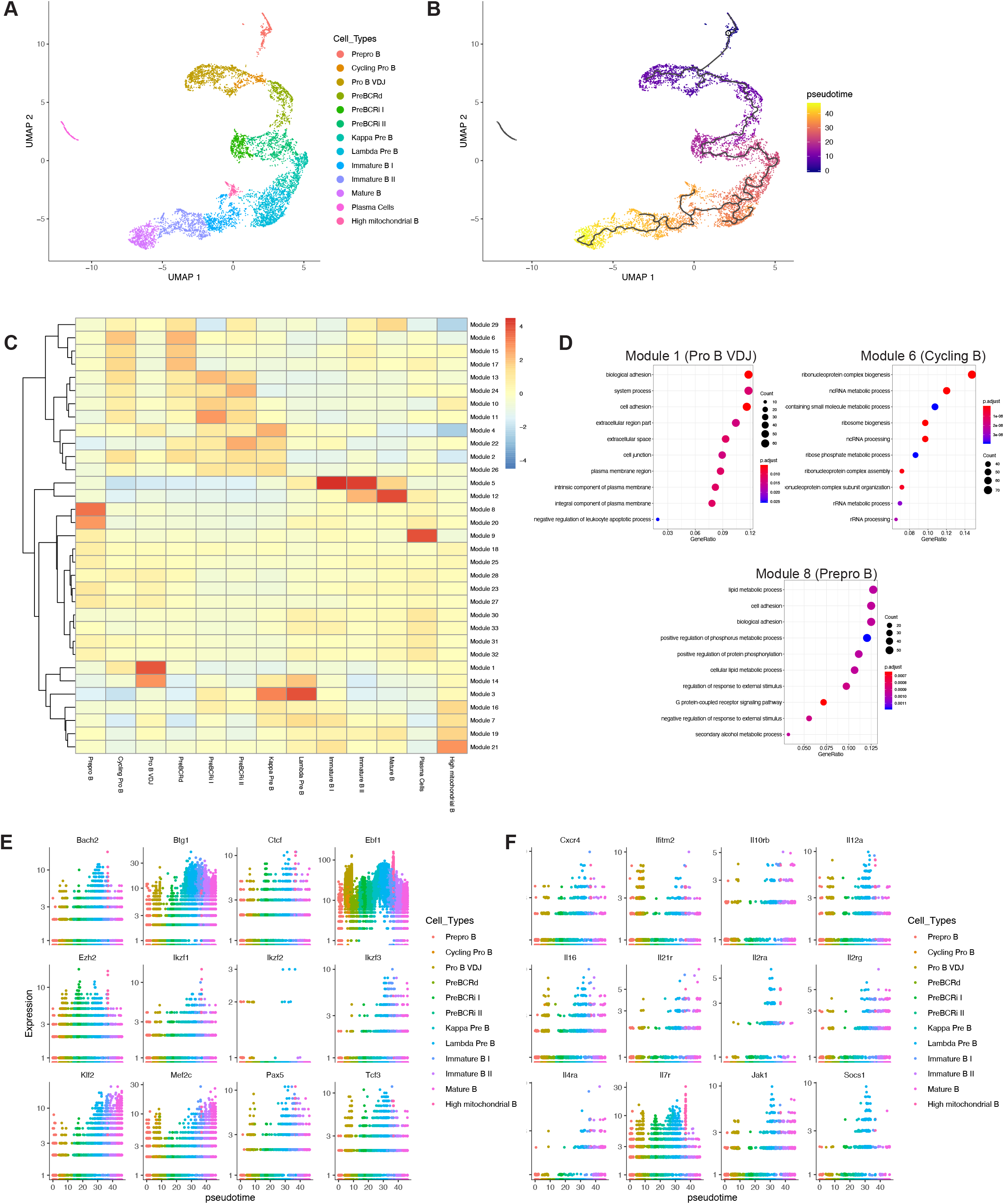
Pseudotime analysis reveals the kinetics of transcription factor expression and gene modules that are differentially expressed across the B cell development trajectory. **A.** Cell cycle-related genes were regressed within Monocle3 and UMAP dimension reduction was performed. **B.** Pseudotime values were calculated and plotted. **C.** Module analysis to demonstrate gene modules that change across the B cell developmental trajectory. A total of 33 modules and their expression intensity for each stage are shown. **D.** Gene ontology term analysis of selected modules 1, 6, and 8. **E.** Expression of B-lymphoid transcription factors and epigenetic factors across B cell development stages. **F.** Expression of cytokine, chemokine and cytokine/chemokine receptors across the B cell developmental trajectory.

Furthermore, the cycling pro-B cells and pre-BCR-dependent cells were enriched for genes involved in metabolic processes (Module 6; Figure 6D), suggesting metabolic reprogramming after major checkpoints (B cell lineage commitment and pre-BCR selection, respectively) during B cell development. Finally, evaluation of the expression of transcription factors, epigenetic modifiers (Figure 6E), cytokines, chemokines and their respective receptors (Figure 6F) revealed differential expression kinetics across the B cell development trajectory. Thus, our findings confirm some previous observations, but also identify distinct gene programs that exhibit highly dynamic changes during B cell development.

### Stage-specific gene expression networks are exploited by human B-ALLs

Defects in B cell differentiation and dysregulation of signaling pathways lead to B cell transformation. To identify B cell developmental pathways that are exploited by human B-ALL subtypes, we used differentially upregulated gene markers identified in both the Seurat and Monocle analyses. The pre-BCR module genes, including *YBX3* and *NRGN*, are significantly upregulated in BCL2/MYC, IKZF1 N159Y, and MEF2D B-ALL subtypes (Figure 7A), which are all associated with high risk and poor prognosis (Gu et al., 2016; Liu et al., 2015; Mullighan et al., 2009). Despite the numerous pre-BCR module genes highly expressed in various B-ALL subtypes, not all pre-BCR genes share this pattern, indicating transcriptional heterogeneity of B-ALLs compared to normal developing B cells. This partial gene expression network usage by various B-ALL subtypes is further appreciated with other B cell development stage markers (Supplementary Figure 6A). For example, the TCF3-PBX1 and MEF2D subtypes have high expression of genes associated with pre-pro B cells and cycling immature B cells (Supplementary Figure 6A). Finally, we examined whether *YBX3* expression correlated with outcome in B-ALL. We observed that pediatric B-ALLs with above-median *YBX3* expression are associated with worse prognosis (Figure 7B; Hazard Ratio = 2.03; p = 0.0315), whereas adult B-ALL patients (which are mainly comprised of Ph^+^ B-ALLs) show no significant difference in survival (Figure 7B). Overall, we identify B cell gene expression networks that are modulated during B cell development (Figure 7C) and are exploited by human B-ALLs. In addition, we specifically demonstrate that a YBX3-related module is associated with a poor prognosis in human B-ALLs.

**Figure 7.**
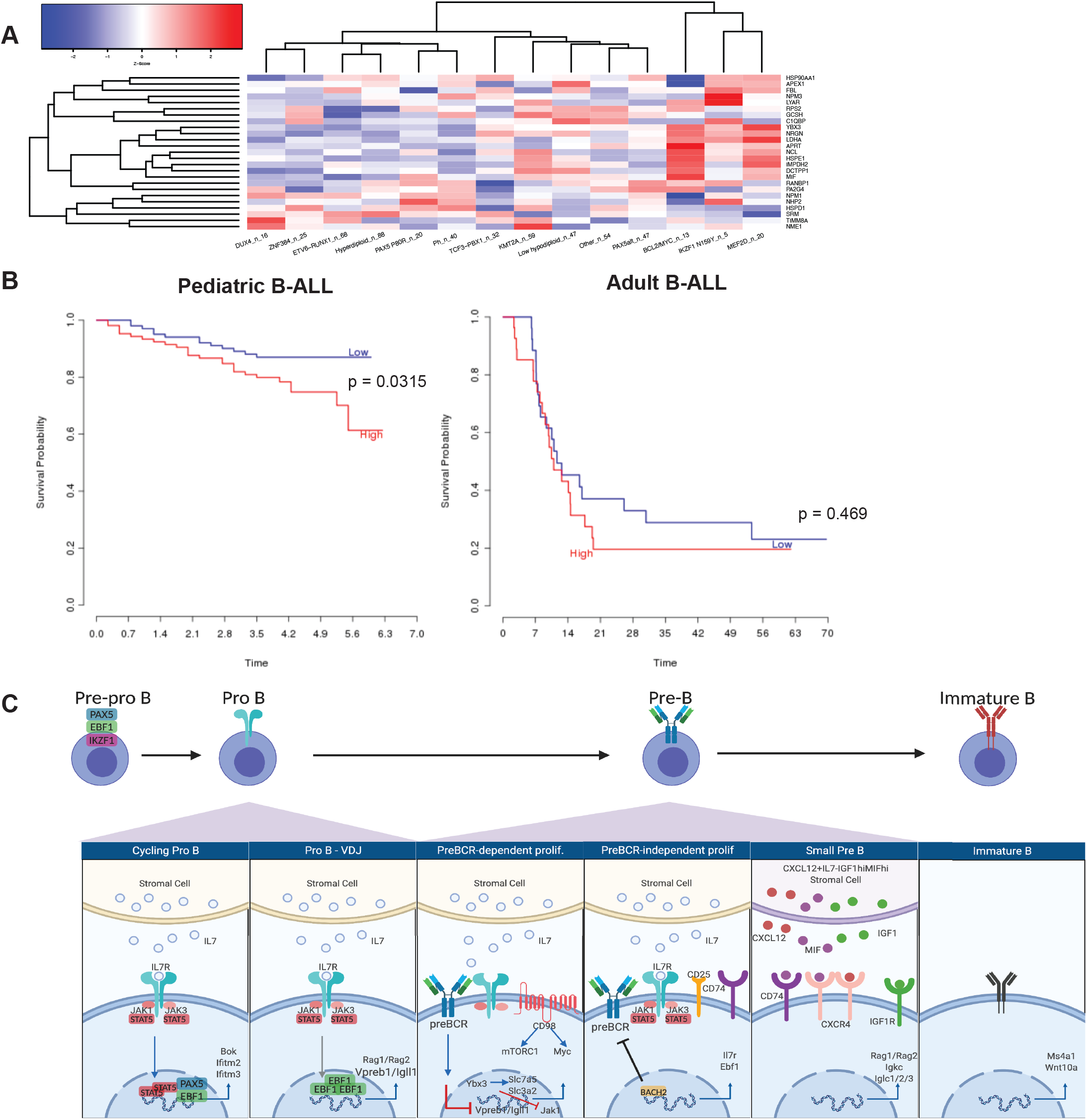
Select B-ALL subtypes exploit pre-BCR-module genes and correlates with poor prognosis. **A.** Overlapping differentially upregulated gene markers in the pre-BCR-dependent clusters from Seurat and Monocle are plotted (y axis). The expression of these genes was queried across various B-ALL subtypes (x-axis). The number of B-ALL samples (n) is listed after each subtype. **B.** Survival curve of pediatric B-ALL (left; time in years) and adult B-ALL (right; time in months). Low and high expression correspond to below median and above median expression of YBX3, respectively. Survival data was obtained from Prediction of Clinical Outcomes from Genomic Profiles (PRECOG)(Gentles et al., 2015) **C.** Proposed model of B cell development generated with Biorender.

## Discussion

The evaluation of different organs (Schaum et al., 2018) and niches (Baccin et al., 2020; Tikhonova et al., 2019) at single cell resolution has greatly expanded our understanding of the cellular diversity that is present within the bone marrow. However, these previous studies have not resulted in a detailed description of B cell development. This is due to the paucity of developing B cells in these broad surveys of the bone marrow compartment. B cell development has also been examined using cell surface markers in conjunction with flow cytometry (Hardy et al., 1991). These studies, in conjunction with in vivo reconstitution experiments to ascertain precursor-progeny relationships, have provided a general outline of B cell development (Clark et al., 2014; Hardy et al., 1991). Furthermore, resources such as Immgen provide a wealth of transcriptional data about various sorted B cell compartments (Heng et al., 2008). However, these sorted cell populations are relatively heterogenous and thus fail to provide detailed single-cell resolution of B cell development. In this study, we coupled the conventional surface marker-based staging of B cell development using CITE-Seq with single-cell transcriptomics to reveal unappreciated transcriptional heterogeneity during B cell development and link them to various underlying biological processes. In addition, we identify the RNA binding protein YBX3 as a novel marker of pre-B cell differentiation and correlate *Ybx3* expression with outcomes in patients with B cell acute lymphoblastic leukemia.

Our studies demonstrate unexpected dynamic changes in transcription factors that play key roles during B cell development. We were largely able to confirm the expression kinetics for the critical B-lymphoid lineage transcription factors *Pax5* and *Ikzf1* across B cell development stages (Figure 3B). In contrast, *Ebf1*, another key B cell transcription factor, has a strikingly large dynamic range of expression (Figure 3B) that varies greatly over the course of B cell development. *Ebf1* has the highest expression during both heavy and light chain recombination stages, intermediate expression during cytokine-mediated proliferative stages, and the lowest expression during pre-BCR and BCR signaling stages (Figure 3B). Likewise, the expression of EBF1 target genes exhibited the same patterns (Figure 3C). This points to a key role for dynamic changes in *Ebf1* in regulating key gene expression networks throughout B cell development. Furthermore, B-lymphoid transcription factors, including *Ebf1*, have been shown to serve as metabolic gatekeepers, where they repress genes encoding proteins for glucose uptake and utilization, and thereby prevent malignant transformation (Chan et al., 2017). In light of this, we provide evidence that EBF1 represses *Myc* and that low gene expression of *Ebf1* correlates strongly with the activation of pre-BCR-dependent gene modules that promote metabolic reprogramming (such as amino acid transporters) and proliferation. Therefore, the expression kinetics of *Ebf1* are tightly controlled to enable the unique aspects of alternating cycles of proliferation and differentiation throughout B cell development.

Our single-cell RNA-Seq data pointed to an unappreciated role for the RNA binding protein YBX3 in pre-B cell differentiation and proliferation. Notably, *Ybx3* expression was significantly upregulated during the pre-BCR dependent stage (Figure 3A). Likewise, the related YBX family member *Ybx1* is also most highly expressed at the pre-BCRd stage (Figure 3A). Analysis of *Ybx3*^*−/−*^ mice did not show major developmental defects (Supplementary Figure 3B and 3C) and thus YBX1 likely serves a redundant function with YBX3 in developing B cells. Therefore, YBX1 and YBX3 may play a critical role in initiating the MYC-dependent transcriptional program that characterizes the pre-BCR-dependent expansion stage (Sinclair et al., 2013).

Elucidating the requirements and regulation of normal B cell development has significantly improved our understanding of how developing B cells can undergo transformation and maintain leukemic states. B-ALL subtypes exhibit significant transcriptional diversity (Gu et al., 2019). Our comparative analysis between stage-defining gene expression in wild-type progenitor B cells and different B-ALL subtypes revealed both overlapping and non-overlapping gene modules, suggesting that B cell leukemias can partially exploit specific gene expression modules of multiple B cell development stages. In particular, we found that pre-BCR module genes, including *Nrgn* and *Ybx3*, were significantly upregulated in the high risk BCL2/MYC, IKZF1 N159Y and MEF2D B-ALL subtypes. In addition to our findings on the importance of MYC during pre-BCR signaling, previous studies have demonstrated the upregulation of IKZF1 and MEF2D (Herglotz et al., 2016) upon pre-BCR signaling. Therefore, aberrant dysregulation through mutations and rearrangements in *IKZF1* and *MEF2D*, respectively, may lead to the heightened usage of pre-BCR-signaling related genes for transformation. Importantly, we were able to demonstrate that high expression of *Ybx3*, a key component of the pre-BCR signaling module, correlates with poor prognosis in pediatric B-ALL (Figure 7B). Interestingly, high expression of *Ybx3* also correlates with dramatic downregulation of *Ebf1*. The combination of elevated YBX3 and reduced *EBF1* (Harvey et al., 2010) expression likely plays a key role in establishing the MYC driven transcriptional program that contributes to pre-B cell transformation. As YBX3 has been shown to repress JAK1 translation (Cooke et al., 2019), this may explain the requirement for ectopic activation of the JAK/STAT5 pathway to overcome a potential YBX3-driven negative feedback loop. Collectively, our data and previously published findings converge on multiple mechanisms that can activate the pre-BCR-signaling gene module leading to high risk leukemia and a poor prognosis. Using strategies that target YBX3 or pre-BCR-related modules in pre-BCRd-related leukemias may improve outcomes in patients with these subsets of high-risk B-ALL.

## Acknowledgments

We thank G. Hubbard, A. Rost, and N. Keller, for mouse and technical assistance, J. Motl, and P. Champoux for cell sorting and Flow Cytometry Core Facility maintenance at University of Minnesota (5P01AI035296), D. George for providing *Ybx3*^*−/−*^ bone marrow, E. Stanley, J. Daniels, and K. Beckman for 10X genomics capture, sequencing and University of Minnesota Genomics Center maintenance, Dr. Meinrad Busslinger for *Pax5*^*+/−*^ mice, Dr. Rudolf Grosschedl for *Ebf1*^*+/−*^ mice, Dr. Tim Ley for *Ybx3*^*−/−*^ mice and J. Pereira, T. Lebien, and D. Owen for review and comments on the manuscript. The authors acknowledge the Minnesota Supercomputing Institute (MSI) at the University of Minnesota for providing resources that contributed to the research results reported within this paper. R.D.L. was supported by an individual predoctoral F30 fellowship from the NIH (F30CA232399). M.A.F. was supported by NIH grants R01AI124512, R01AI147540, R01CA232317.

## Author Contributions

Conceptualization, R.D.L. and M.A.F.; Methodology, R.D.L., and M.A.F.; Formal Analysis, R.D.L., S.A.M., T.P.K., R.S.L., L.H.H., M.A.F.; Investigation, R.D.L, L.H.H.; Resources, M.A.F.; Data Curation, S.A.M., T.P.K., R.S.L.; Writing – Original Draft, R.D.L., M.A.F.; Writing – Review & Editing, R.D.L, S.A.M., T.P.K., R.S.L., M.A.F.; Visualization, R.D.L.; L.H.H.; Supervision, M.A.F.; Project Administration, M.A.F.; Funding Acquisition, R.D.L, M.A.F.

## Methods

### Animals

All animals used were bred and housed in specific pathogen-free facilities at the University of Minnesota and Washington University in St. Louis and animal experiment protocols were approved by Institutional Animal Care and Use Committees. All of the animals used were 6- to 10-week old C57BL/6J males and females with appropriate age- and sex-matched controls. The *Ybx3*^*−/−*^ mice were graciously provided by Dr. Timothy Ley at Washington University in St. Louis and have been previously described (Lu et al., 2006).

### Tissue processing and cell preparation

For flow cytometry and cell sorting experiments, bilateral femurs and tibias were harvested from mice. Bones were flushed with 1X PBS with 2% fetal bovine serum (FBS), 0.1% sodium azide and 0.5 mM ethylenediaminetetraacetic acids, pH 7.4. The cells were filtered through a 70 μm mesh, centrifuged at 1200 rpm for 5 minutes and then incubated for 5 minutes with 5 mL of ACK lysis buffer for red blood cell lysis. Cells were then washed and centrifuged at 1200 rpm and subsequently resuspended for cell counting on a hemocytometer (Fisher Scientific) and staining.

### Flow Cytometry and Antibodies

All flow cytometry was performed using the BD Fortessa cytometers (BD Biosciences) in the University of Minnesota Flow Cytometry Core. Bone marrow cells obtained using the method above were stained with various FACS antibodies, including B220-BUV395, CD11c-APCef780, GhostRed780, Ter119-APCef780, NK1.1-APCef780, GR1-APCef780, CD4-APCef780, CD8-APCef780, and CD43-Biotin. In short, surface staining was performed for 20 minutes with FACS antibodies on ice, washed and either analyzed or stained for intracellular FACS antibodies. For intracellular staining of VPREB1-PE (BioLegend), JAK1-AF488 (R&D Systems) and EBF1-PE (BD Biosciences), surface stained cells were fixed/permeabilized using the eBioscience Transcription Factor staining kit (eBioscience) for 30 minutes at room temperature, washed, then stained for 30 minutes in permeabilization buffer. Cells were then washed and resuspended in 1X PBS with 2% fetal bovine serum (FBS), 0.1% sodium azide and 0.5 mM ethylenediaminetetraacetic acids, pH 7.4 for flow cytometric analysis. Cell sorting was performed on a BD FACSAria sorter (BD Biosciences). All flow cytometry data acquired was analyzed using FlowJo software (Tree Star)

### Cell Hashing and CITE-Seq

Bone marrow cells from two 8-week-old wild-type mice were each stained with 1 μg of different hashtag antibodies (TotalSeq A0301 and TotalSeqA0302, respectively; BioLegend). At the same time, cells were stained with 1 μg of FACS antibodies: B220-Pacific Blue (BioLegend), CD43-Biotin (BD Biosciences), CD11c-APCef780 (eBioscience), Ter119-APCef780 (eBioscience), NK1.1-APCef780 (eBioscience), Gr1-APCef780 (eBioscience), CD4-APCef780 (eBioscience), CD8-APCef780 (eBioscience), GhostDye Red780 viability dye (Tonbo Bioscienes) and 1 μg of CITE-Seq antibodies (BioLegend): B220 (TotalSeq-A0103), CD19 (TotalSeq A0093), CD93 (TotalSeq A0113), CD25 (TotalSeq A0097) and IgM (TotalSeq A0450). Cells were stained with the above-mentioned antibodies for 20 minutes on ice, washed and resuspended in 1X PBS with 2% fetal bovine serum (FBS), 2 mM ethylenediaminetetraacetic acids, pH 7.4 buffer containing 1 μg of streptavidin-PE (TotalSeq A0113), which served the dual purpose of cell sorting and CITE-Seq for the CD43 antigen expression. scRNAseq was performed in parallel to FACS analysis.

### 10X Genomics 3’ v3 Single-cell RNA-sequencing

For 10X Genomics scRNA-seq, we generated three libraries that measure: (1) mRNA transcript expression (RNA), (2) mouse-specific hashtag oligos (HTO), and (3) cell surface marker levels using antibody derived tags (ADT). Cells were harvested and stained as described above. In order to have balanced populations of various B cell development stages, we enriched for earlier progenitor B cells (B220^+^CD43+) by sorting at a 1:1 ratio of Dump^−^B220^+^CD43^+^ cells and Dump^−^B220^+^CD43^−^ cells. A total of 20,000 cells per mouse were sorted into a single microtube containing 1X PBS with 50% FBS and were washed and resuspended in 1X PBS with 10% FBS prior to cell capture. The sample was split into three libraries (RNA, HTO, and ADT). Reverse transcription PCR and library preparation were carried out under the Chromium Single Cell 3’ v3 protocol (10X Genomics) per manufacturer’s recommendations. After library prep, quality control was performed using a bioanalyzer (Agilent 2100 Bioanalyzer; Agilent Technologies) and preliminary sequencing of the RNA library on a MiSeq (Illumina) to determine the approximate number of cells and general quality. After passing quality control, the library was sequenced on the NovaSeq 6000 with 2×150 bp paired-end reads (Illumina). Raw and processed data has been deposited at Gene Expression Omnibus and is available via GEO accession #. The code used in this study can be obtained upon request.

### Single-cell Bioinformatic Analyses

Raw sequencing data was processed using the CellRanger pipeline (version 3.1.0; 10X Genomics) “mkfastq”to demultiplex the three Illumina libraries (RNA, HTO, and ADT) and “count” was used to align reads to the mouse genome (mm10, provided by 10X Genomics, ver 3.0.0) and generate mRNA transcript, HTO, and ADT count tables. Raw count data were loaded into R (v. 3.6.1) and analyzed with the Seurat R package (v 3.0.3.9039). The RNA dataset was filtered to include only GEMs (droplets with uniquely barcoded beads that ideally contain one individual cell) expressing more than 300 genes (counts > 0) and genes expressed in more than 3 GEMs (counts > 0). The proportion of mitochondrial RNA in each GEM was calculated and GEMs with extreme levels (top 0.5% of all GEMs) were removed from the analysis. For the remaining GEMs, the HTO count table was added to the dataset, normalized by a centered-log ratio method, and used to determine whether the GEM contained a single cell or multiple cells. The Seurat function “HTODemux” was used to classify what cells were detected in each GEM using the expression of the HTOs as markers. The function was tested using a range of initial k-values (7 – 26), where k = 22 provided the cleanest classification results. A total of 7,454 GEMs contained WT singlets (3,902 WT-1 or 3,552 WT-2) and 966 GEMs contained multiplets (removed from further analysis). The WT singlets expressed a median of 1,409 genes with a median of 3,548 counts. For the WT singlets, the raw RNA counts were transformed using the Seurat function “SCTransform” (Hafemeister and Satija, 2019) including the percent of mitochondria expression as a regression factor. Principal components analysis (PCA) was performed using the normalized SCT dataset (RunPCA function) and two-dimensional representations were generated using the top 30 PCA vectors as input to the RunTSNE and RunUMAP functions. Cells were clustered using the FindNeighbors function (top 30 PCA vectors) and FindClusters function (testing a range of possible resolutions: 0.05, 0.1, 0.15, 0.2, 0.25, 0.3, 0.4). A final resolution of 0.4 was used to classify cells into gene expression clusters. Each cell was classified according to its expression of canonical cell cycle genes using the CellCycleScoring function. The SCT normalized expression values were used for calculating S and G2M phase-specific scores (genesets provided in Seurat). These S and G2M phase scores for each cell were used as additive factors in a linear model of gene expression (i.e. when regressing out the influence of cell cycle).

Single cell surface protein expression data (ADT) was filtered to include only the WT singlets and counts were normalized according to the centered-log-ratio method in Seurat. For each of the six markers measured (B220, CD19, CD93, CD25, IgM, CD43), the normalized counts were centered (subtracting the mean expression from each value) and scaled (dividing centered value by standard deviation) across all cells. The Seurat object with S and G2M phase scores and a resolution = 0.4 was converted into a cell_data_set object for use with the Monocle (v3) R package(Cao et al., 2019). The aligncds function was used with residual_model_formula_str = “~S.Score + G2M.Score” to adjust for the cell cycle status. After adjustment for cell cycle status, UMAP dimensional reduction and clustering were performed in Monocle. The final resolution for Monocle clustering was 0.0009. This resolution resulted in 3 separate partitions for the clusters. The plasma cell cluster was in its own partition and was excluded from the Pseudotime trajectory analysis. The remaining two partitions were (1) pre-pro B cells and (2) all other cell populations. We relabeled the partitions so that the pre-pro B cells and the remaining cell populations of interest would be in the same partition for pseudotime analysis. We initiated the pseudotime trajectory in the pre-pro B cell population, using a custom function described in the Monocle 3 documentation (get_earliest_principal_node) to automatically select the starting point for the pseudotime analysis.

We used Monocle 3 methods to identify modules of co-regulated genes. First, we used the function graph_test to identify variable genes in the data using the Moran’s I statistic. Then we identified the genes that had a significant q-value (< 0.05) from the autocorrelation analysis and then grouped these genes into modules using UMAP and Louvain community analysis. We used the enrichGO function in the clusterProfiler package (v 3.14.3) to evaluate enrichment of the modules in GO terms across all three ontologies (BP, CC, and MF)(Carbon et al., 2019; Yu et al., 2012).

### Landscape In silico Deletion Analysis (LISA)

The top 100 differentially upregulated genes obtained from Seurat FindAllMarkers function was used as input into the LISA Cistrome (lisa.cistrome.org). The transcription factor ChIP-Seq dataset was used to infer the transcriptional regulators for differentially regulated genes of the pre-BCR-dependent proliferation stage.

### Enrichment Analysis of Single Cell RNA-Seq Cluster Marker Genes in Human B-ALL Gene Expression

We used enrichment analysis to examine whether Human B-ALL subtypes have similarity to particular B cell progenitor populations. To create robust signatures for each cell type from the single cell data we used the intersection of the Seurat and Monocle top cluster marker genes for each of the different cell types to create our combined cluster marker gene sets. The top marker genes are the genes that are differentially expressed for a given cluster when compared with all other clusters. Default approaches for finding top marker genes were used for both Seurat and Monocle. We downloaded publicly available B-ALL bulk RNA-Seq count data(Gu et al., 2019). We summed expression values for each gene across biological replicates for each B-ALL subtype to create an average sample for each sub-type. We normalized and log2-transformed the data to create log2cpm values for unsupervised hierarchical clustering of each combined cluster marker gene set against all B-ALL subtypes.

#### Statistical Analysis

Differential gene expression (DE) analysis was completed using the FindMarkers function, employing a Wilcoxon rank sum test between all pairwise clusters or between a single cluster vs. all others. Genes were considered significant if the absolute value of log2-fold-change was >= 0.25 and Bonferroni adjusted p-value <= 0.01. Data and statistical analyses were performed using Prism 8 (Graphpad). A Shapiro-Wilk test was performed to assess data normality, and unpaired data that passed normality was analyzed using an unpaired student t-test.

## Supplemental Information

**Supplementary Figure 1.** Both wildtype samples are well-represented in all clusters. **A.** UMAP plots showing cells colored based on origin of wildtype sample (left) or colored by cluster assignment and separated according to each wildtype sample (right). **B.** ADT/CITE-Seq antibody expression in wildtype sample #1 cells. **C.** ADT/CITE-Seq antibody expression in wildtype sample #2 cells.

**Supplementary Figure 2.** Pre-B cell expansion gene expression. **A.** Violin Plot gene expression of Pre-BCR-dependent stage genes (*Nrgn*, *Ybx1*, *Ybx3*, *Slc7a5*, *Slc3a2*, *Myc*). **B.** *MKi67* expression Violin Plot for all B cell development stages.

**Supplementary Figure 3.** Flow cytometric characterization of the role of YBX3. **A.** Flow cytometry analysis of WT developing B cells in the bone marrow gated on B220^+^ and CD98^high^ cells (left). B220^+^CD19^+^ CD43^+^CD98^high^ cells were gated and displayed for KI67 and JAK1 expression (middle). JAK1 expression was split into low and high subsets and EBF1 expression analyzed (right). **B.** Flow cytometric analysis of B cell development from wildtype and *Ybx3*^*−/−*^ mice pregated on B220^+^CD43^+^ progenitor B cells and stained for BP1 and CD24 expression (Hardy Fractions A-C). **C.** Flow cytometric analysis of B cell development from wildtype and *Ybx3*^*−/−*^ mice pregated on B220^+^CD43^−^ progenitor B cells and stained for B220 and IgM expression (Hardy Fractions D-F). **D.** Summarized percentage of lymphocytes for each Hardy fraction (Fraction A-F) between wildtype and YBX3^−/−^ B cells. Unpaired student t-test was performed. Error bars indicate standard deviation. Flow plots represent n=3 experiments and data are represented as mean ± S.D.

**Supplementary Figure 4.** Genes and surface protein expressions that are differentially expressed during B cell differentiation and maturation. **A.** Feature Plot of selenoprotein genes **B.** IgM surface expression across B cell development stages.

**Supplementary Figure 5.** Validation of cell clusters after cell-cycle regression based on stage-specific gene expression. **A.** Feature plots for various stage-specific genes to characterize the clusters after cell cycle gene regression

**Supplementary Figure 6.** Different gene expression modules from multiple stages of B cell development are exploited by various human B-ALL subtypes. **A.** Overlapping differentially upregulated gene markers from pre-pro B, cycling pro B, pro B VDJ, pre-BCRi I, Kappa PreB, Cycling Immature B, Mature B and Plasma cells are plotted. Gene expression intensity for each of these genes in different B-ALL subtypes is shown as a heatmap.

**Supplementary Table 1.** Differential gene markers of every cluster compared to all clusters. Seruat was used to identify the differential genes expressed of every cluster compared to all clusters. Avg_logFC represents log_2_ fold change

**Supplementary Table 2.** Differentially expressed genes identified from pairwise comparison of each cluster. Seruat was used to identify the differential genes expressed from a pairwise comparison of each cluster. Avg_logFC represents log_2_ fold change

