## Supplemental figures for "Single-cell analysis of developing B cells reveals dynamic gene expression networks that govern B cell development and transformation"

**Supplementary Table 2.** Differentially expressed genes identified from pairwise comparison of each cluster. Seruat was used to identify the differential genes expressed from a pairwise comparison of each cluster. Avg_logFC represents $\log_2$ fold change

Supplementary Figure 1

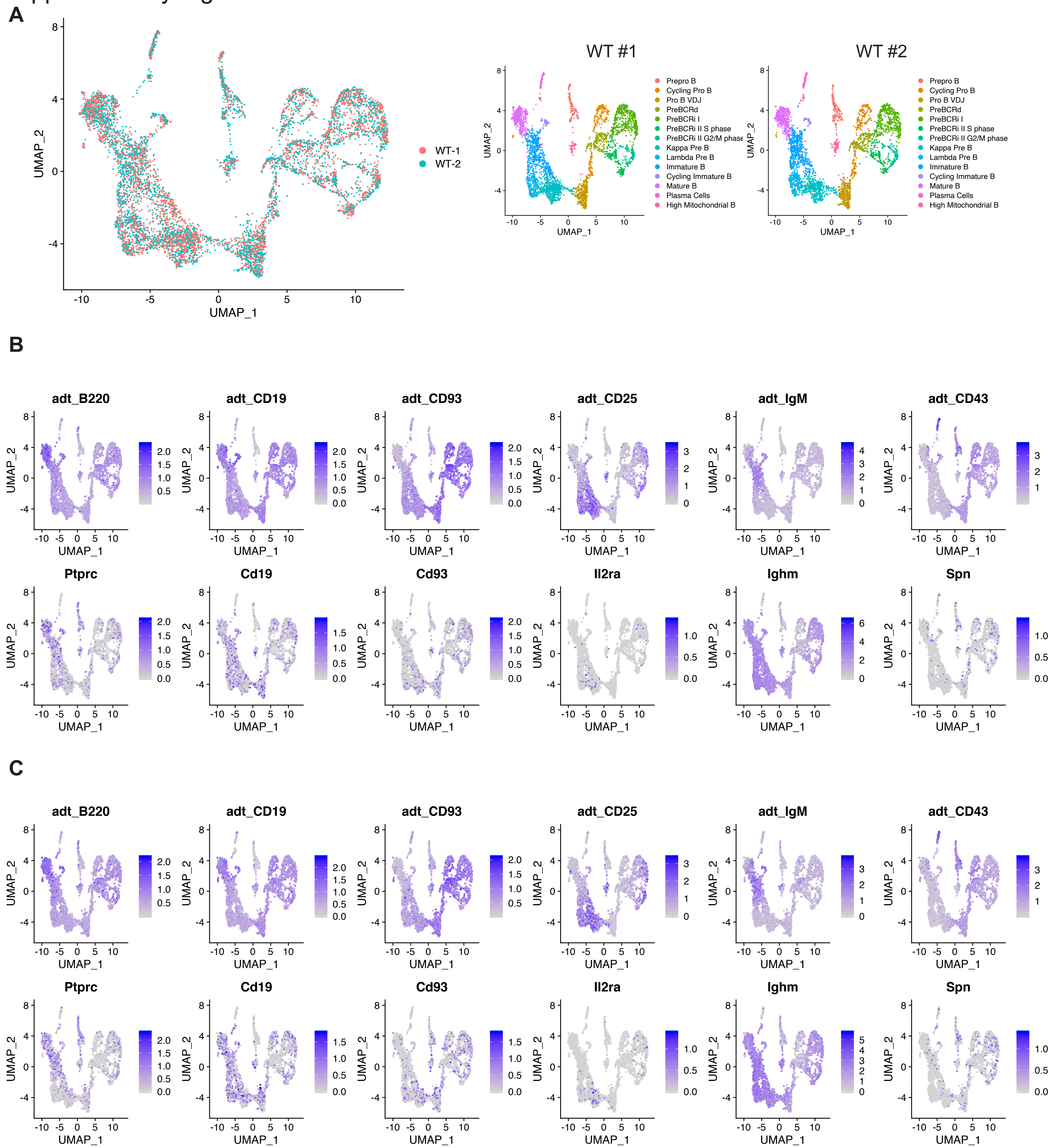

### Supplementary Figure 2

**A**

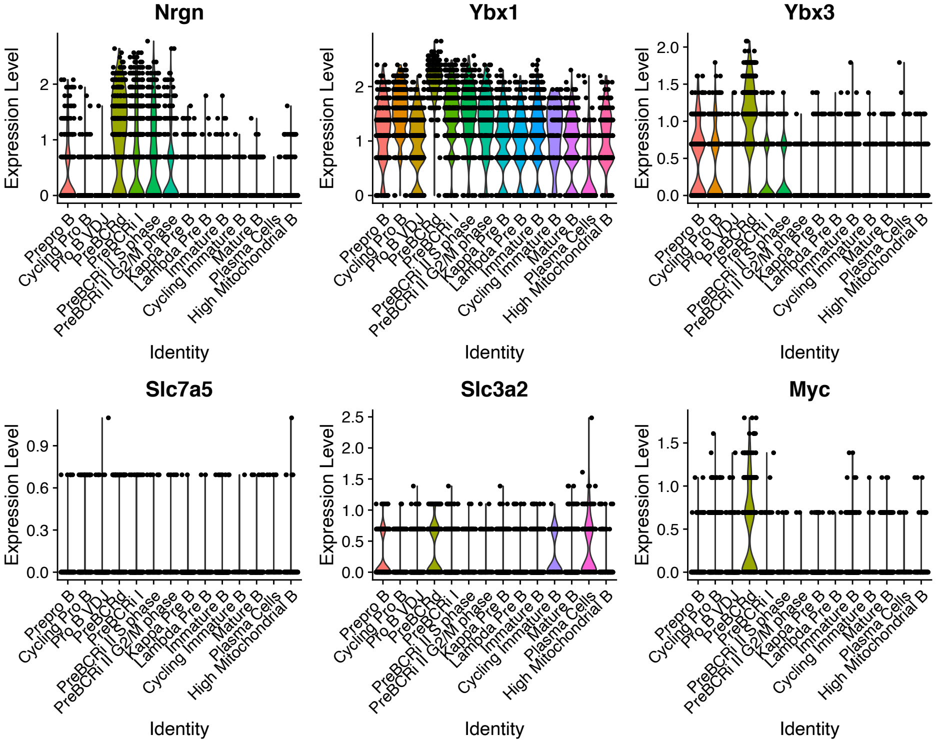


**B**

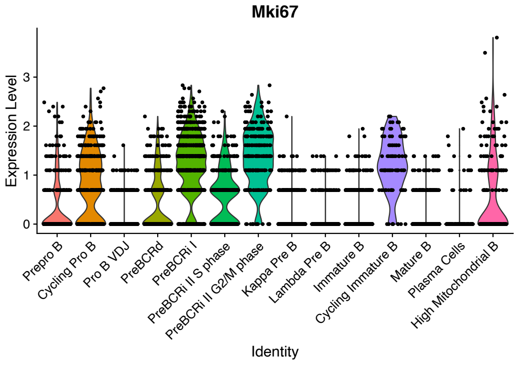

Supplementary Figure 3

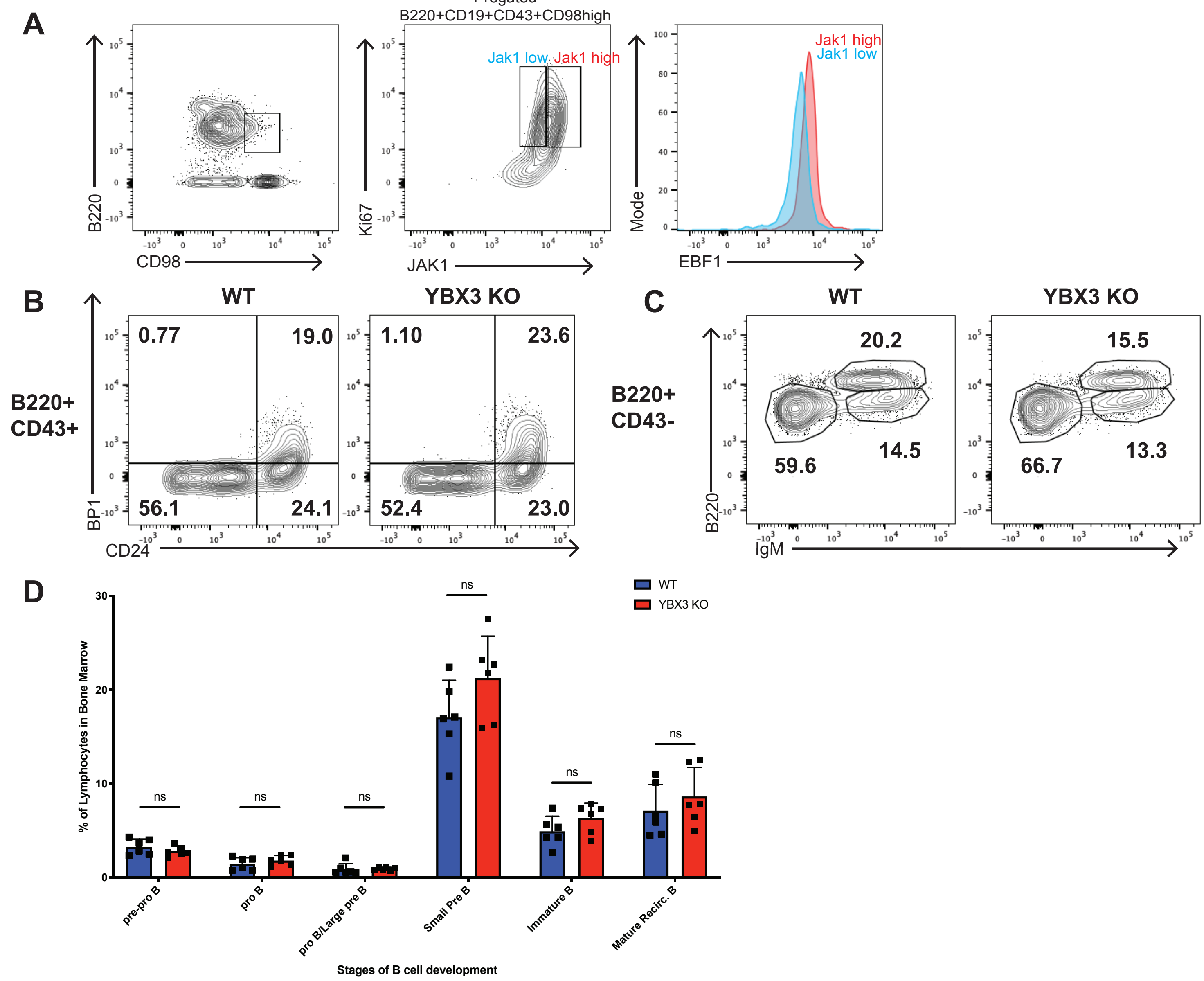

Supplementary Figure 4

**A**

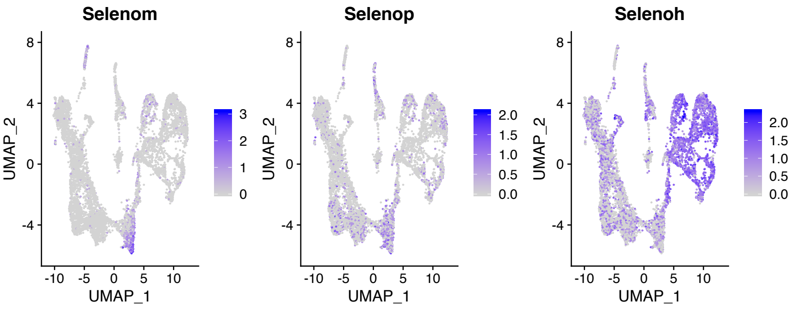


**B**

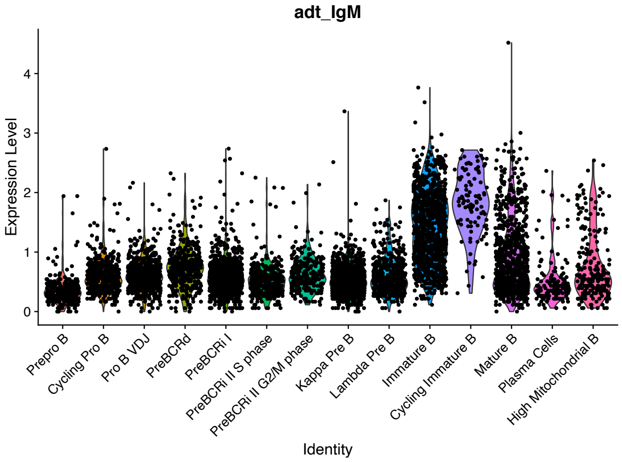

### Supplementary Figure 5

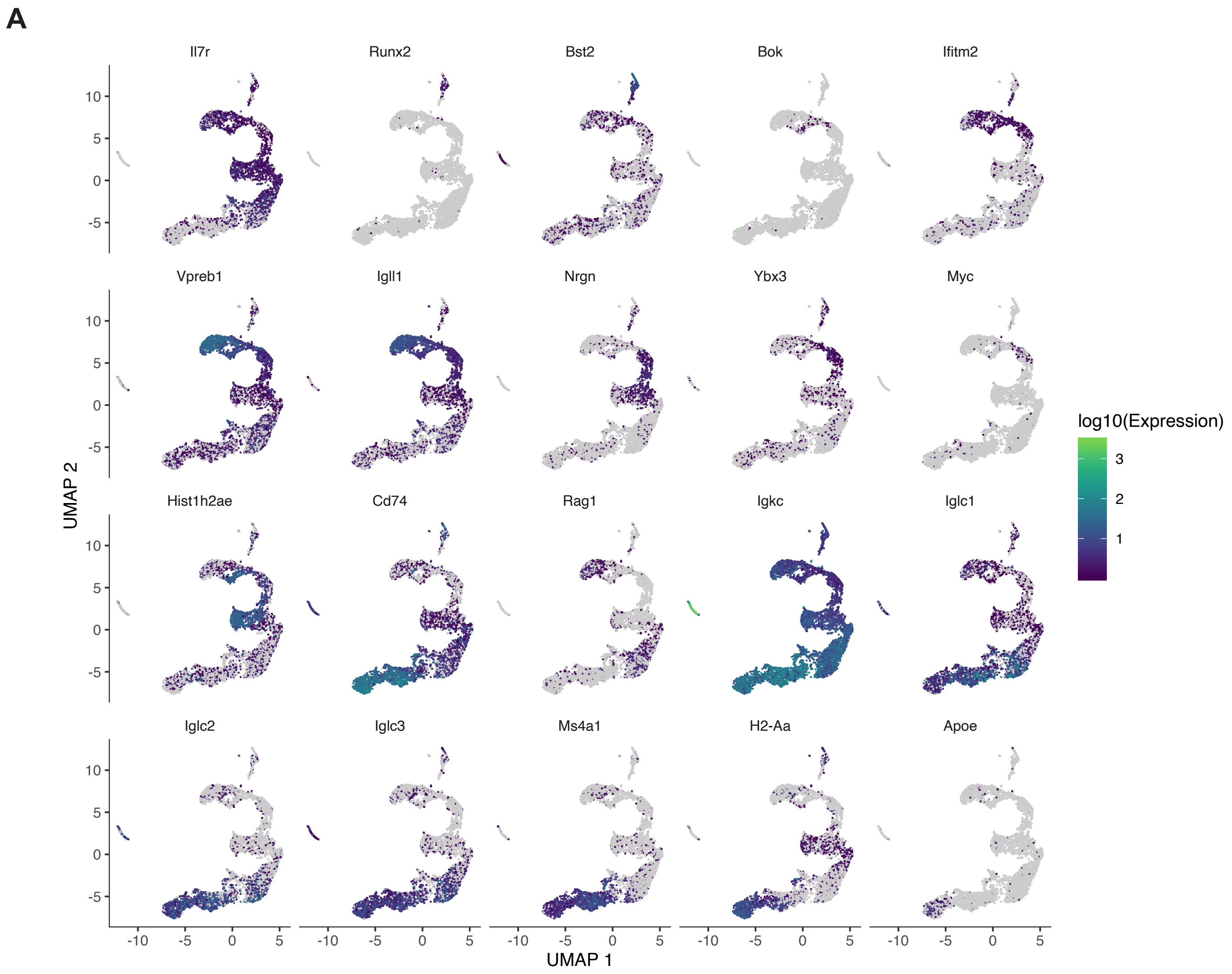

### Supplementary Figure 6

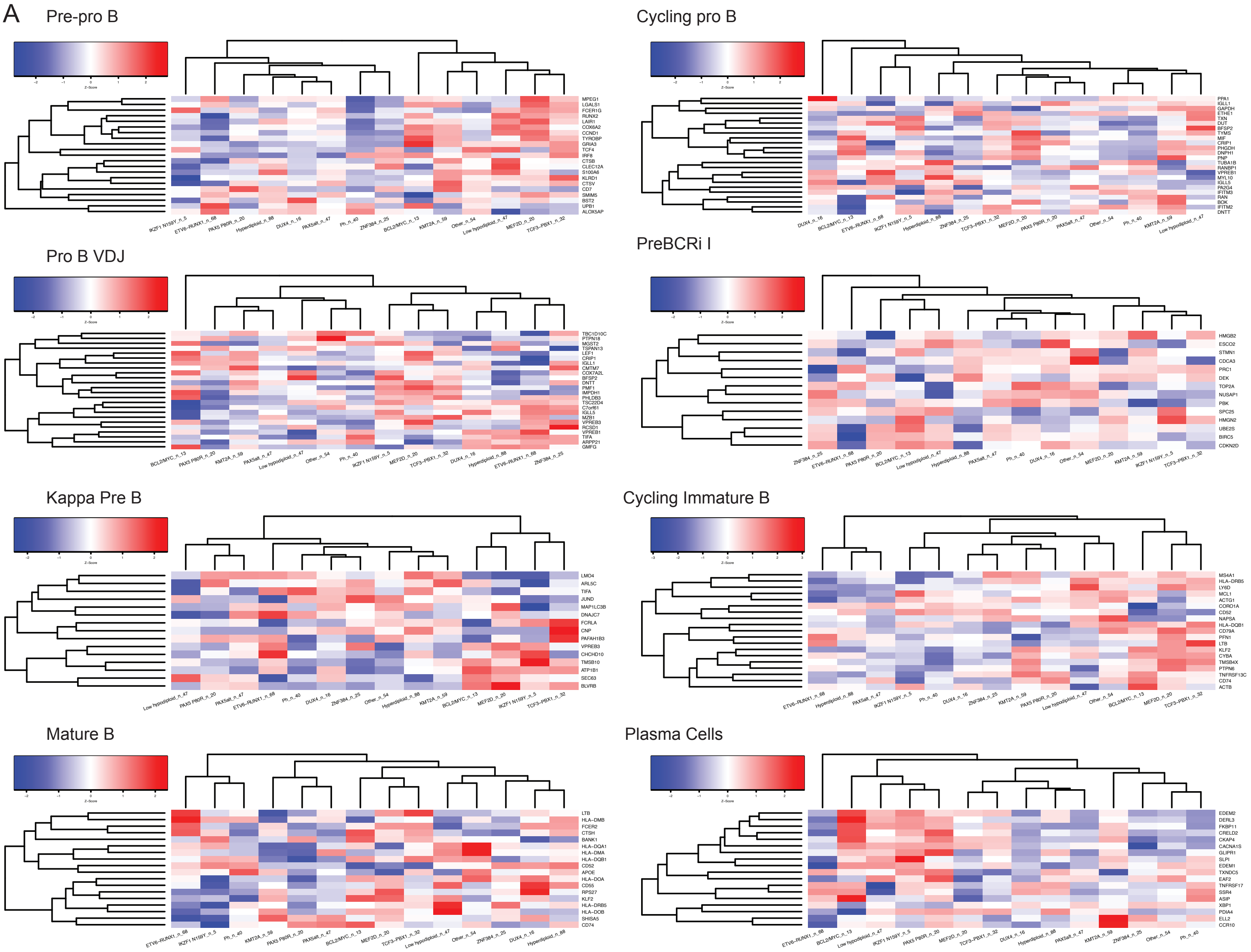
